# CCSeq: Clusters of Colocalized Sequences

**DOI:** 10.1101/818385

**Authors:** Stefan Golas

## Abstract

**Motivation:** Potential transcription factor (TF) complexes may be identified by testing whether the binding sequences of individual TF proteins form clusters with each other. These clusters may also indicate TF inhibition due to competitive occupancy of enhancer regions. Genome annotation data containing the coordinates of enhancer sequences is highly accessible via position-weight matrix tools.

**Results:** An algorithm called CCSeq (Clusters of Colocalized Sequences) was developed for identifying clusters of sequences along a one-dimensional line, such as a chromosome, given genome annotation files and a cut-off distance as inputs. The algorithm was applied to the binding sequences of the constituent proteins of two known transcription factor complexes, the HSF1 homotrimer and one form of the NF-*κ*B complex, a dimer of NFKB2 and RELB. 28 clusters of HSF1 trimer binding sequences were identified on chromosome Y, and 16 clusters of the NFKB2 and RELB dimer were identified on chromosome 17, compared to 0 clusters identified in any of the five simulated random distributions for each of the two sets of TF proteins. Additionally, structural patterns of these binding sequence clusters are described.

**Availability and Implementation:** This algorithm is freely available as an R package on the open source R repository CRAN at the following link: https://cran.r-project.org/package=colocalized. Genome annotation files were obtained from the PWMScan tool at https://ccg.epfl.ch/pwmtools/pwmscan.php hosted by the Swiss Insitute of Bioinformatics (2) (3).

## 1 Introduction

Transcription factor (TF) proteins are a key element in gene regulatory circuits and other mechanisms for controlling gene expression (7), and many diseases are related to mutations in TFs. Many TFs bind with each other to form complexes, such as the heat shock factor 1 (HSF1) homotrimer (9) or the NF-*κ*B heterodimer (6). These and other complexes are critical regulators of cell responses to stimuli, and have been implicated in a wide range of diseases including cancer. It is reasonable to assume that the different subunits of a TF complex will have DNA-binding sequences in close physical proximity to each other along a chromosome, and we can exploit this tendency to identify potential complexes using only sequence data. An algorithm CCSeq was created in which the user provides an input selection of sequence sets, where each set corresponds to the binding sequences of a particular TF (these binding sequences are also known as enhancers). The algorithm identifies clusters of DNA-binding sequences colocalized to each other within a given cut-off distance. The first step of the algorithm identifies pairs of colocalized binding sequences across sequence sets, each set corresponding to the binding sequences of a transcription factor in the input selection, and the locations of these sequence pairs are stored in colocalization matrices. Next, pairs of colocalized sequences are compared across multiple colocalization matrices to identify chains of paired sequences. Very simply, if we find that a sequence *a* pairs with sequence *b* and *b* pairs with *c*, the cluster *abc* is recorded. A cluster will be defined as a set of sequences where every member is colocalized to at least one other member of the cluster, two sequences are colocalized if they are within a given cut-off distance of each other. A number of papers have studied combinatorial binding in TFs, but these either rely on overlapping binding regions from individual ChIP-Seq experiments or otherwise take into account other measurements such as chromatin accessibility (8), whereas this algorithm relies solely on genome annotation files generated via position-weight matrices. The terms sequence and interval are used somewhat interchangeably in this paper. For clarity, a sequence is a substring along a chromosome whose interval is its starting and ending coordinates. Since this paper is concerned with the spacing and distribution of sequences rather than their contents, the interval will suffice to refer to the sequence and vice versa.

## 2 Algorithm

The input selection given to the algorithm consists of multiple sets of DNA-binding sequences, where each sequence set corresponding to a particular TF protein. A cut-off distance for colocalized pairs is also provided in the input. The two main steps in the algorithm are the creation of colocalization matrices for each pair of sequence sets, and the multiplication of these matrices in dimensionally sound sequences to locate clusters. The details of these steps will be elaborated in section 2.2, while an illustration of a simple example is given in figure 2.2.

### 2.1 Notation

Upper-case letters will in general refer to matrices or sequence sets, and lower-case letters to integers. The colon “:” will index all the elements of a particular row or column of a matrix, e.g. *K*_:,*z*_ will refer to all elements in the *z*th column in *K*. Certain variables *i,j,k*, and *z* are used to represent different values at different points throughout the pseudocode where they are used to index arbitrary elements in a vector or matrix. The meaning of each usage of these variables should be clear within each context.

### 2.2 Pseudocode

#### Algorithm 1 CCSeq

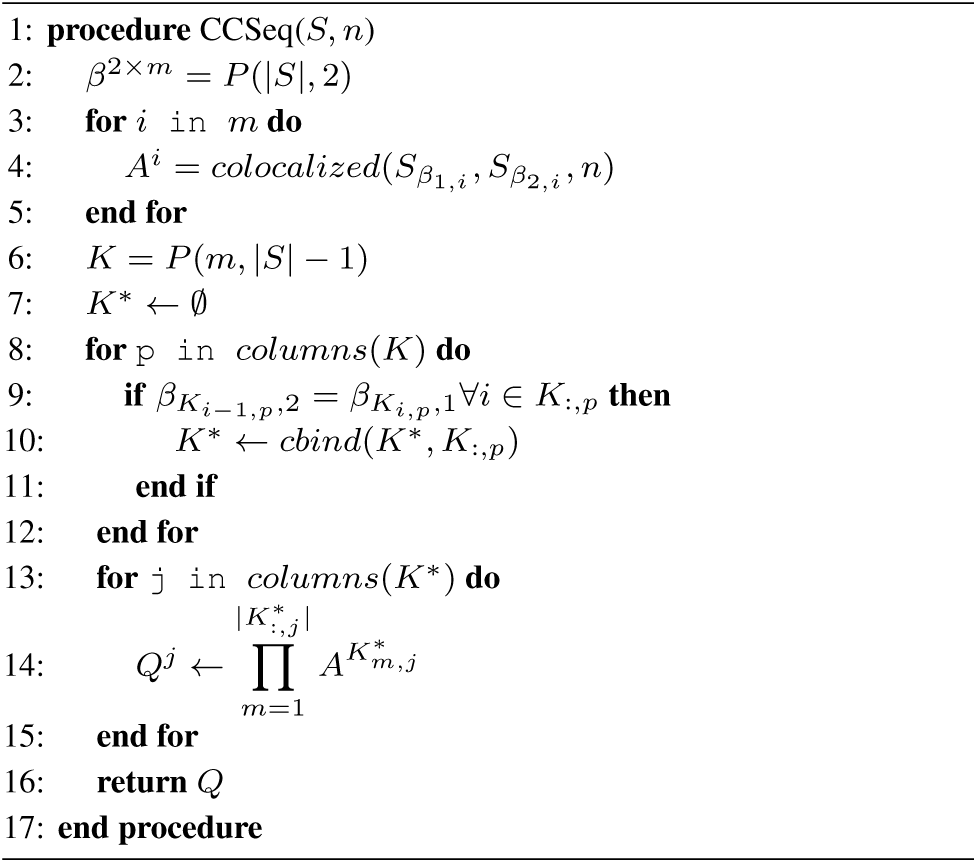

For an input selection S and a cut-off distance n, the function CCSeq(S,n) proceeds according to the pseudocode. Line 2 generates all 2-element permutations on the index of S and stores these in the matrix *β*. Lines 3-5 iterates the function *colocalized*(*X, Y, n*) over columns of *β*^2*×m*^. Very simply, *colocalized*(*X, Y, n*) forms a colocalization matrix such as has been already described from two sequence sets *X* and *Y* and a cut-off distance. For a colocalization matrix *M* = *colocalized*(*X, Y, n*), a matrix entry *M*_*i,j*_ = 1 where the *i*th element of *X* and the *j*th element of *Y* are less than *n* base pairs from each other and 0 otherwise. In lines 6-12, a matrix *K* is formed containing every permutation over the index of *A* containing |*S*| − 1 elements with permutations stored as columns. These columns are filtered to form the matrix *K\**, which will index elements of *A* = *A*_*i*_, *A*_*j*_, … such that the sequence set represented by the column of some 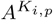 is the same as that represented by the row of 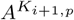. This will ensure that taking the product of some sequence of colocalization matrices in *A* indexed by a column in *K\** will be logical according to the formula for forming a cluster by chaining colocalization pairs with shared partners. If the columns of a colocalization matrix *A* represents the same sequence set as that represented by the rows of a matrix *B*, then *C* = *AB* will have entries *C*_*i,k*_ = *A*_*i*,1_*B*_1,*k*_ + *A*_*i*,2_*B*_2,*k*_… + *A*_*i,j*_ *B*_*j,k*_ where the shared sequence set has length *j*. The formula for entries of the composite matrix *C* formed from *A* and *B* attempts to form clusters by finding where colocalized pairs from *A* and *B* share a sequence from the matching sequence set, and an entry *C*_*i,k*_ = 1 where *A*_*i,j*_ = 1 and *B*_*j,k*_ = 1. This can be considered in terms of Boolean logic, where *C*_*i,k*_ is *true* if there exists a *j* where *A*_*i,j*_ and *B*_*j,k*_ are both *true*, and the summation to find *C*_*i,k*_ checks every possible value of *j*. In this sense Boolean logic can entirely replace the standard arithmetic formula and the same result will be arrived at. Indeed, this fact is exploited in the implementation of this algorithm for significantly improved run-time. Lines 13-15 perform this matrix multiplication in series on colocalization matrices in *A*. The loop which filters *K* to *K\**uses the functions *columns*(*K*) and *cbind*(*K*, K*_:,*p*_). The function *columns*(*K*) extracts from *K* the index over its columns, such that an element *j* from *columns*(*K*) can be used to extract a column from *K* via *K*_:,*p*_ where the “:” symbol implies every entry in the column *p*. The function *cbind*() creates a matrix by binding the columns of two input matrices. It is here used exactly as it is used in the language R. After performing the sequence matrix multiplication in lines 13-15, matrices *Q*^*j*^ ∈ *Q* will contain a positive entry 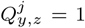 where colocalization pairs had shared partners at every matrix in the matrix multiplication sequence. For example, if *Q* = *ABC*, then *Q*_*y,z*_ = 1 where (*AB*)_*y,y″*_ *C*_*y″, z*_ = 1 for some *y″* and *A*_*y,y′*_ *B*_*y′, y″*_ = 1 for some *y′* and the same *y″*. Therefore, positive values in *Q*^*j*^ and its factors will index the locations of individual sequences belonging to clusters within their respective sequence sets.

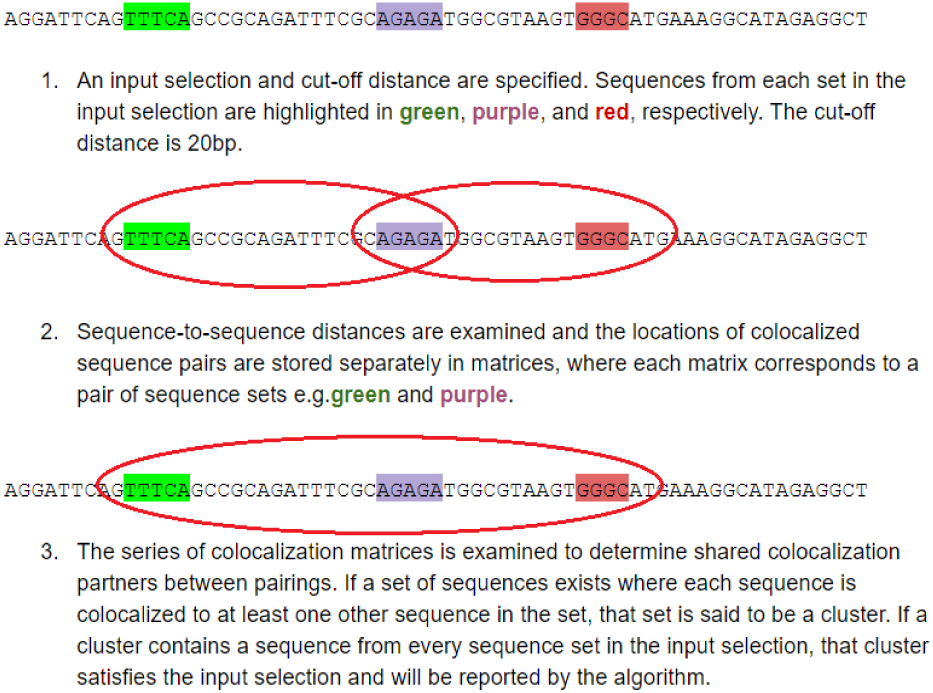

## 3 Implementation

Two known TF complexes, the HSF1 homotrimer and the NF-kB heterodimer (composed of NFKB2 and RELB), were tested as inputs to the algorithm on chromosome Y and chromosome 17, respectively, each with cut-off distances of 150bp. 16 clusters of NF-kB dimer sequences were detected on chromosome 17, while 28 clusters of HSF1 trimer binding sequences were detected on chromosome Y. Further data on these clusters are given in the supplementary data. Not only are clusters of binding sequences associated with the formation of complexes by proteins, the structure of the clusters in the two example sets is notably specific. In the case of the NF-kB complex, NFKB2 and RELB bind to complementary sequences on opposite strands, so the presence of one binding sequence almost always implies the presence of the other. For the HSF1 homotrimer, the three binding sequences are always on the same strand; 16*/*28 clusters are spaced according to the pattern [115, 30] and 5/28 are spaced according to the pattern [99,69], with the remaining 7/28 not conforming to any regular spacing pattern. Spacing patterns are denoted according to [x,y..] where x is the distance between the lowest-numbered sequence and the 2nd-lowest-numbered, and so forth.

### 3.1 Simulating a Null Hypothesis

It is necessary to consider that these clusters may exist by chance and that their existence or frequency of occurrence has no significance as to whether the proteins of their constituent sequences form a binding complex. For a null hypothesis, we assume that individual binding sequences are distributed randomly across the specified chromosome. If there is a sequence set *x* in the input selection, then to test the null hypothesis a sequence set called *x*^′^ is generated with the same number of sequences as *x* and whose intervals are all the same size as that of the average interval size in *x*, but which are distributed randomly throughout the length of the corresponding chromosome. A randomized sequence set is generated for every member of the input selection, and the algorithm is executed on the randomized selection with the same parameters (i.e. cut-off distance and chromosome). In order to reject the null hypothesis, the algorithm must detect consistently more clusters in the input selection than in the randomized sequence sets. In the examples of HSF1 and NF-kB given above, no clusters were detected in randomized sequence sets across five randomized samples each. In this case the null hypothesis is rejected.

## 4 Discussion

Combinatorial regulation of gene expression has been a focus of intense study for years, in part due to its role as a built-in platform for cellular logic circuits and computation (18) (19) (20) (21) (22) (23). It has long been understood that these logic circuits can be exploited for metabolic engineering of industrially relevant cell lines (25) (26) (27) (28). Additionally, networks of interacting TFs play a key role in defining cancer cell lines (29). Clustered enhancer sequences within the genome provide crucial evidence for characterizing transcriptional circuitry and networks, and when combined with patterns of gene co-expression and protein-protein interaction data, may be used to defintiively identify candidates for TF complexes. Although this paper has primarily focused on TF complexes as the motivation for identifying clustered binding sequences, transcriptional repression of one TF by another via competitive occupancy of enhancers is another potential explanation for clustering with experimental evidence (30). The structure of binding-sequence clusters may also be used to investigate patterns in the physical interaction of TF proteins with each other and the chromatin itself. The strength of this algorithm in investigating elements of transcriptional circuitry is in the high accessibility of genome annotation data given the availability of web tools such as PWMScan and JASPAR.

### 4.1 Comparison with other algorithms

Although other algorithms and studies have examined combinatorial TF regulation involving proximal binding sites (10) (11) (12) (13), this algorithm diverges from those in several ways. In general other algorithms investigating clusters of enhancers for different TFs have relied on overlaps between ChIP-Seq peaks, which recover accessible binding sites on chromatin for specific cell types and experimental conditions. This is because ChIP (chromatin immunoprecipitation) studies require that a physical binding event take place in order to detect a signal, which depends highly upon chromatin accessibility due to epigenetic regulation, which itself depends on complex external and internal signals (14) (15) (16) (17). This makes ChIP-Seq highly appropriate for investigationg the interaction of environment, epigenetics, and cell fate, but it will fail to identify enhancer clusters in regions that are effectively “switched off” under the conditions of a given experiment. CCSeq relies exclusively on genome annotation files generated by the application of PWMs to whole genomes, and will identify clusters in locations which may not be accessible to TFs under a given set of experimental conditions, or any conditions whatsoever. From this perspective, ChIP-Seq based algorithms should be used as part of a targeted approach under specific experimental conditions, while CCSeq offers a more exploratory approach that can be applied to genome annotation files generated using widely available web programs and open-source data. The algorithm will also report the exact locations of clustered binding sequences rather than regions with overlapping peaks.

## Supporting information

Supplemental table NFKB clusters

Supplemental table HSF1 clusters

## 5 Acknowledgements

Thank you to Ed Reitman and Jack Tuszynski for supporting this project through its duration with your excellent feedback and ideas. Thank you Philip Winter, Jack Tuszynski, and the University of Alberta for providing access to the Compute Canada research cluster which was invaluable for testing this algorithm and developing its parallel computing capabilities. Thank you to Renato Spacek for helping with the math very early in the project.

